# Unraveling the Role of G-Quadruplexes in Alternative Polyadenylation (APA): A Focus on Neogenin 1

**DOI:** 10.1101/2024.06.04.597383

**Authors:** Pauline Lejault, François Bolduc, Marc-Antoine Turcotte, Jean Pierre Perreault

## Abstract

Polyadenylation is a crucial step in mRNA maturation. In half of the cases, alternative polyadenylation sites are used instead of canonical ones, affecting the length of the transcript’s 3’UTR. The mechanisms controlling the selection of these alternative polyadenylation sites are still unknown. Using a unique sequencing method (PolyAclick-seq) to identify different polyadenylated isoforms, we selected events modulated by RHPS4, a ligand known to stabilize G-quadruplex (G4). Through *in silico* selection, *in vitro* assays, and G4 mutagenesis construction, the pivotal role of rG4 structures in determining polyadenylation sites was uncovered, particularly for the gene encoding Neogenin-1 (NEO1). This research highlights the importance of focusing on G4 RNA mediated APA regulation in the 3’UTR, as a method to alter isoform choice and impact protein synthesis opening up new avenues for RNA-based therapies.

## Introduction

Advancements in RNA biology have opened new avenues in RNA-based therapeutics such as mRNA vaccines and antisense oligonucleotides.^1^ The development of therapeutics requires a deeper understanding of RNA structures, and its interaction with host molecules in cells, particularly how to modulate mRNA stability as to induce a prolonged and more efficient expression and response. In Eukaryotes, pre-messenger RNA (pre-mRNA) maturation requires splicing and polyadenylation. Polyadenylation, the addition of a polyadenine tail at the 3’ of pre-mRNA, is essential for mRNA export, stability and translation. ^2^ Numerous pre-mRNA contains multiple PolyAdenylation Sites (PASs), yielding alternative isoforms with different PolyA tails. This phenomenon, known as Alternative PolyAdenylation (APA), is widespread, affecting 50 to 70% of transcripts, and significantly shapes the transcriptome (through 3’UTR length) and proteome (via altered coding sequences), impacting RNA/protein levels, subcellular localization, and even protein functions. ^3,4^ Unlike splicing, APA site recognition proceeds sequentially and appears to be independent of whether it’s the distal or proximal isoform. While dysregulation of APA has been linked to numerous pathological conditions such as cancer, neuropathologies, and cardiac dysregulations, the precise mechanisms driving these dysfunctions remain elusive. ^5,6^

rG4s are RNA guanosine-rich sequences that fold into non-canonical 4-stranded helical structures. Depending on their localization, rG4s are suspected to fine tune pre-mRNA and mature mRNA lifecycle. For instance, in the 5’UTR, they affect the accessibility of the mRNA to the translation machinery resulting in either promotion or inhibition of translation. ^7,8^ In pre-mRNA, they can modulate alternative splicing events by acting as splicing regulatory elements, thus impacting the inclusion or exclusion of specific exons in the mature mRNA. ^9^ In 2013, our lab unexpectedly discovered the involvement of RNA G-quadruplex (rG4) in APA within the 3’-UTR of two transcripts (i.e., LRP5 and FXR1). rG4s structures located at 49 and 60 nucleotides from APA sites, respectively, can influence the selection of APA sites, leading to shorter alternative transcripts and can interfere with miRNA regulatory networks.^10^ Despite the critical importance of APA in pre-mRNA stability and functional protein production, the formation of secondary structures and their implications remain largely unknown to date.

In this study, we explore the role of rG4s in APA at the transcriptome level through the employment of the innovative PolyA click-seq methodology. Strikingly, our analyses revealed that the small molecular tool, RHPS4, also known as a G4-ligand, induces significant alterations in the PolyA profile within cells. Among these, our investigation into Neogenin-1 revealed that the formation of the 3’UTR distal rG4 seemed to exert a stronger influence on the selection of PAS site and the production of long isoforms in comparison to the 3’UTR proximal rG4. Our findings carry significant implications for mRNA maturation and protein production. Previous research, including our own, has shown that therapeutic targeting of the 3’UTR can alter protein production. Here, we unveil a novel layer of APA regulation involving G4-RNA. Our data suggest that targeting G4-RNA-mediated APA regulation could hold promise as a therapeutic strategy for several previously mentioned disorders.

### A novel dataset of APA induced by RHPS4

In order to highlight the role of potential G4s (pG4) as a universal regulator of APA, we used a well-established G4-ligand, 3,11-Difluoro-6,8,13-trimethylquino[4,3,2-kl] acridinium methylsulfate (RHPS4), in HEK-293T cells.^11^ RHPS4 has previously been identified as a potent inducer of alterations in the G4-RNA landscape across both coding and non-coding mRNAs, leading us to anticipate modifications in the APA landscape.^12–14^ Initially, we determined by MTT assay the inhibitory concentration of RHPS4 to evaluate the suitable concentration for use on HEK-293T cells (Supp. Fig. 1). Based on the results, the cells were treated at the IC_20_ (1.5 μM) of RHPS4 for 72 h. Following treatment, total RNA was isolated and libraries were prepared to conduct PolyA Cluster-seq (PAC-Seq), aiming to identify significant alterations at the APA level and modification in gene expression (Fig. 1).^15,16^ These innovative approaches include both the use of computational tools to re-analyze RNA-seq to identify APA events as well as targeted sequencing approaches customized to focus on the 3′-end of mRNA by click-chemistry. As described previously, raw data have been analyzed using Differential PolyA-Clustering (DPAC) for the preprocessing of polyA-tail targeted RNAseq data. This pipeline includes mapping polyA sites, clustering and annotating them, and determining the usage of differential polyA-clusters through DESeq2.^17^ It has been used to simultaneously measure changes in gene expression as well as detecting APA events with high confidence. The conclusive omics analysis from PAC-Seq enabled us to identify major changes related to the addition of RHPS4:

**Figure 1:**
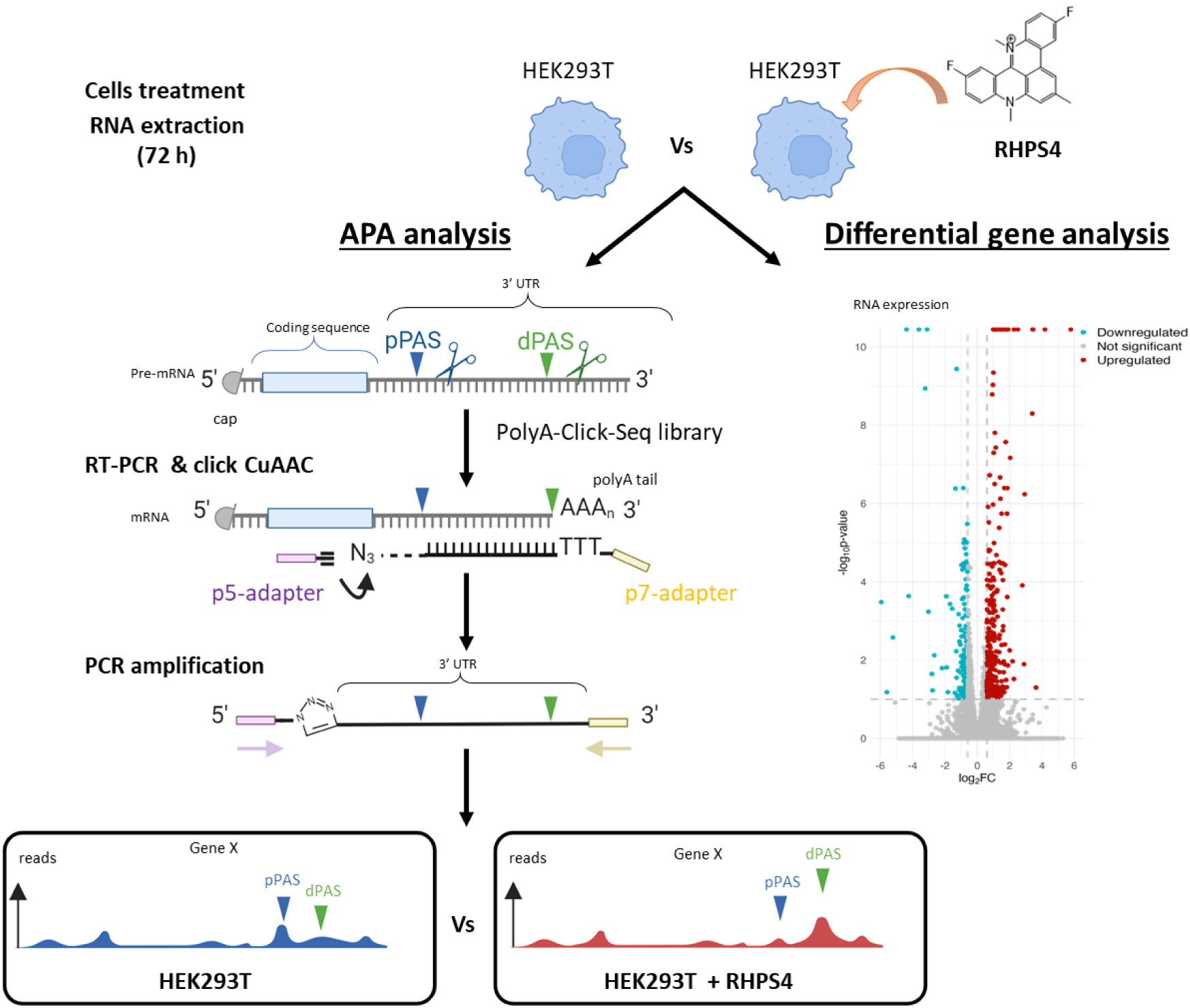
Exploring RHPS4-induced alterations in alternative polyadenylation events: a comprehensive guide in HEK-293T cells. Schematic overview of PolyAClick-Seq (PAC-seq). After several key steps, including RT-PCR, Click-chemistry, and PCR amplification, the final library consists of DNA fragments containing the Illumina p5 adaptor, a portion of the UTR, a stretch of adenine derived from both the RNA template and the poly(T) primer, and finally, the p7 Illumina Indexing primer. Following PAC-seq, the used of DESeq2 analysis allows for the determination of mRNA expression levels, while DPAC software analysis can differentiate between various APA events, such as APA-exon, APA-intron, Splicing, and Multiple exons with PACS.

i) <u>The global RNA differential expression</u>: We identified 14 819 genes including 498 with significant variation in expression (>1.5 fold having a p-adjust <0.1). We noted significant up-regulation variations for 304 genes and down-regulation variations for 194 genes between RHPS4 treated cells and untreated control (Figure 2A). Both lists of up-regulated and down-regulated genes, along with their corresponding GeneOntology (GO) annotations, are presented in Supp. Fig. 2 and Figure 3. The top up-genes are involved in the stress response and the regulation of apoptosis signalisation;

**Figure 2:**
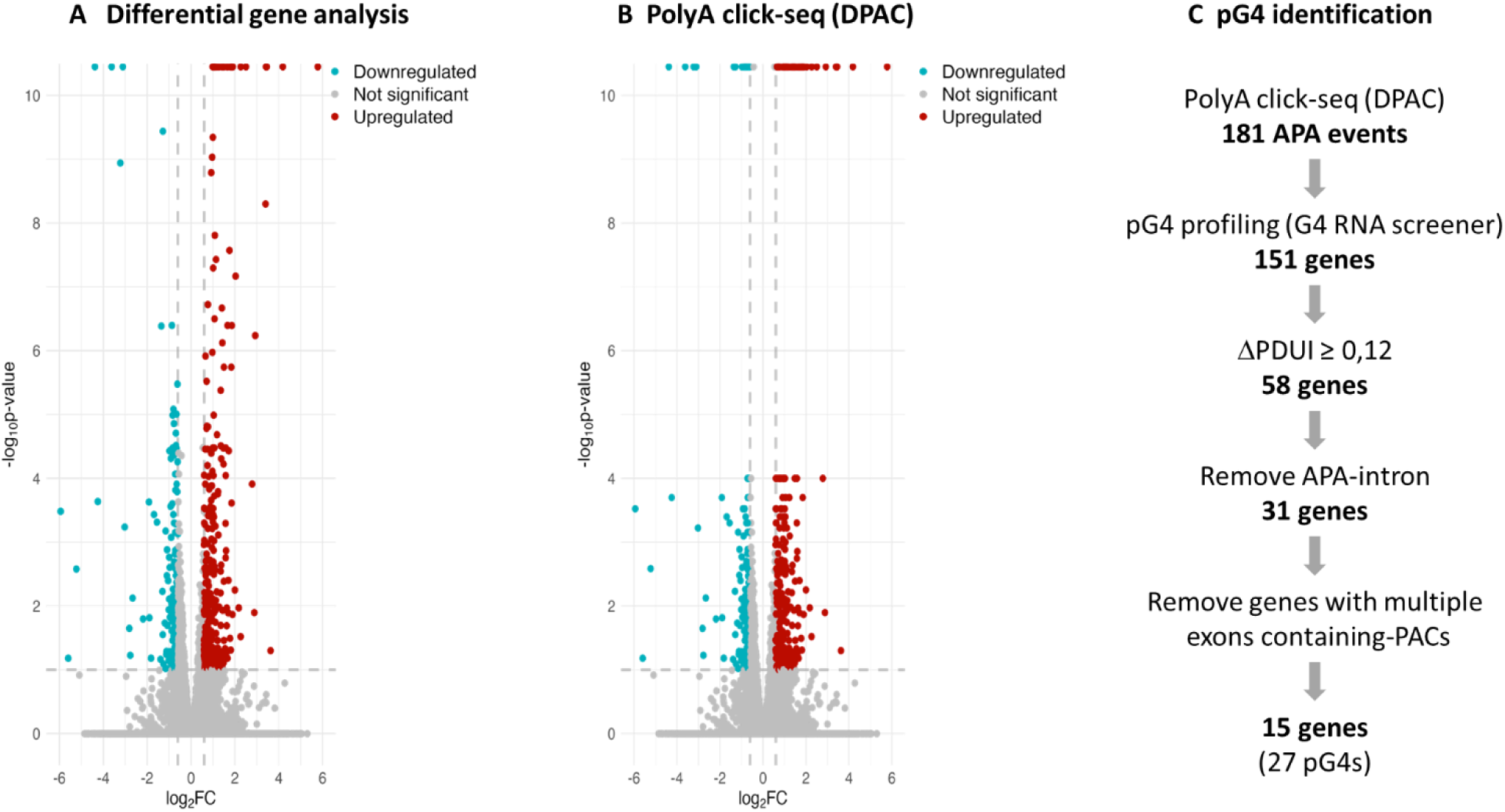
Characterizing changes induced by RHPS4 treatment: impact on gene expression and polyadenylation site usage. (A) Output visualization of DESeq2 results with Log2 fold change versus the P-adjusted value for genes detected in PAC-seq. Genes with a fold change greater than 1.5 and an adjusted p-value less than 0.1 are colored red for upregulated genes and blue for downregulated genes. (B) DPAC results: any alternative polyadenylation changes at the 3’ UTR end with a magnitude exceeding 10% and a p-value <0.1 have been reported. Significant downregulation is indicated in blue, while significant upregulation is indicated in red (C) bioinformatic pipeline for G4 detection: Starting with the 181 genes (DPAC with significant variation = lengthening, shortening or both), a search using G4 RNA Screener for the presence of potential G4s was carried out 100 nt upstream or downstream of the genomic position of the identified polyA clusters (PACs). Subsequently, only ΔPDUI greater than 0.12 are retained. All situations containing more than one exon or involving an intron were also removed.

**Figure 3:**
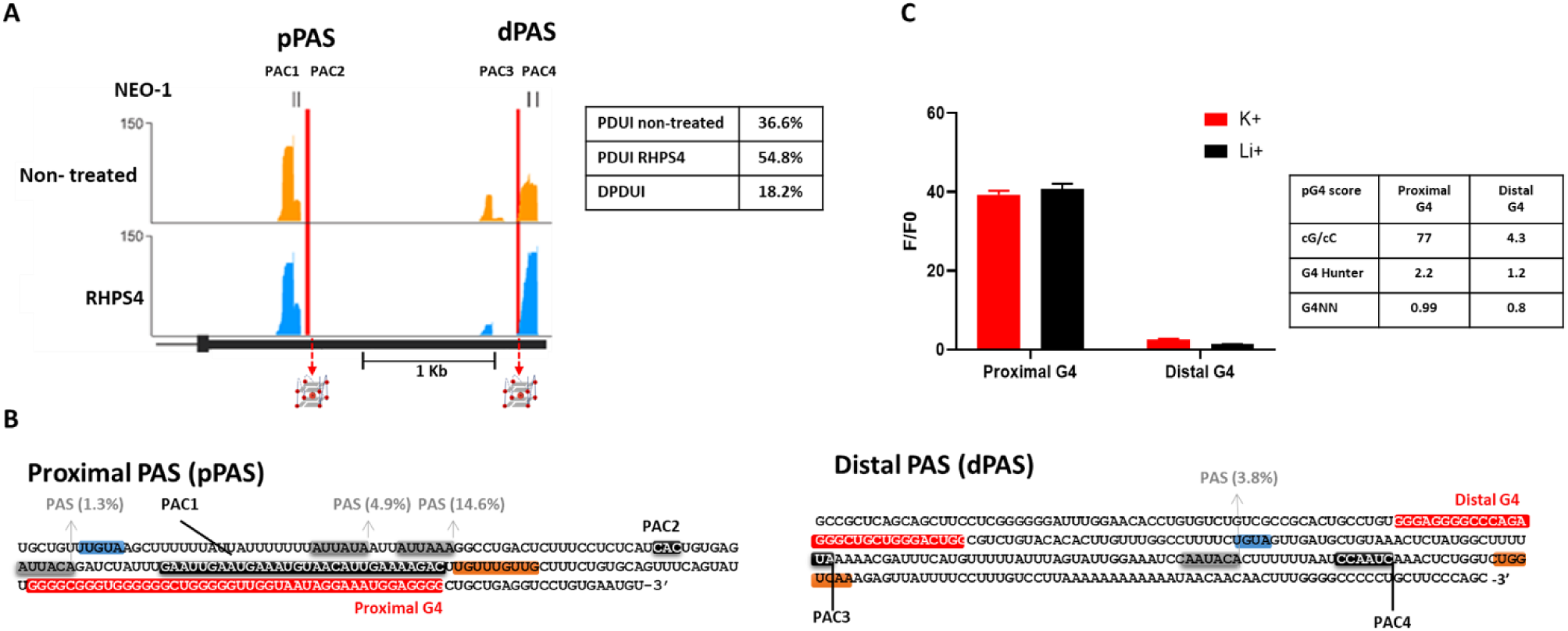
*In vitro* analysis of the potential G4 sequences found in the surrounding area of the proximal and distal polyA sites. (A) Genomic view of the mapped reads found in non-treated and in RHPS4-treated HEK-293T cells in NEO1 3’UTR. The polyA clusters (PACs) are identified by bars at the upper part of the graph (numbered 1 to 4). The red vertical lines represent the position of PG4. On the right, a table with PDUI obtained for NEO1 from DPAC for each condition. (B) The sequence details of the proximal PAS and the distal PAS are as follows: *cis*-elements are highlighted. The polyA Signals site and their percentages of used are shaded in gray, ^29^ the polyA cleavage sites are marked in black, the UGUA motif is indicated in blue, the G/ G-U rich elements (called DSE elements) are highlighted in orange, and the potential G-quadruplex region is depicted in red. (C) Fluorescence assays were performed on *in vitro* transcripts of pG4 derived from both the proximal and distal regions of the NEO1 3’UTR, under conditions with either K^+^ or Li^+^. The histogram illustrates the fluorescence ratio of the G4-light-up fluorophore (NMM) in the presence of RNA compared to NMM alone (F/F0). Furthermore, the analysis includes the pG4 formation, which integrates three scores from the G4 RNA screener.

ii) <u>The significant change in APA</u>: As expected, RHPS4 ligand treatment caused several changes in APA landscape as showed in Figure 2B. A total of 297 significant changes were identified, including both the up-regulated and down-regulated events (see Supp. Table 1). These changes were characterized by the differential usage pattern of a single PAC, as indicated by an Independent Hypothesis Weighting (IHW) p-adjust <0.1, resulting in a fractional change of PAC usage of at least 10%;

iii) <u>The localization of each APA event,</u> (i.e., APA-exon, APA-intron, Splicing and Multiple exons with PACS ^2,17–19^) :When two or more PACs (each with a minimum occupancy of 5% per PAC) are found within a single exon, a change in the abundance of one of the middle PACs can lead to the APA-induced shortening, lengthening, or both in the 3’UTR.Among the APA events induced by RHPS4, 86 events were classified as “splicing,” 36 as “shortening,” 86 as “lengthening,” and 59 as having a dual effect, termed “both” (Supp. Tables 2 and 3); and,

iv) <u>The quantitative point of view of each APA event:</u> Difference in APA usage between normal and treated cells can be quantified as a change in Percentage of Distal polyA site Usage Index (PDUI), which is capable of identifying lengthening (positive index) or shortening (negative index) of 3′ UTRs. (Supp. Table 1). Out of the 27 548 exons with a PAC detected (in 14 819 genes), 37% exhibited a change in PDUI per exon following RHPS4 treatment.

The widespread presence of G4 structures in nucleic acid, suggests that treatment with G4-ligands, such as RHPS4, modulate RNA expression, as demonstrated in previous studies with MCF7 cells.^12^ Here, we present novel findings indicating that a G4-ligand can induce notable changes in the APA profile for the first time in HEK293T. The global impact of G4 stabilization using RHPS4 underscores the potential significance of G4 structures in regulating APA processes.

### Potential G-quadruplex forming sequences are found near PACs with APA

In the entire transcriptome, more than 1 million pG4 sequences are suspected to modulate a plethora of mechanisms, including translation, splicing, and polyadenylation.^20^ The use of RHPS4 has a global impact on G4 formation. Here rG4s involved in APA were selected to conduct more specific *in vitro* experiments. To do so, we conducted a search near the PACs genomic positions (100 nt both upstream and downstream of the PAC) to identify the presence of these structures using G4 RNA screener. ^21^ This was aimed to highlight the role of G4 structures in the APA changes induced by RHPS4. The G4 RNA screener is a bioinformatics tool designed for identifying regions with a probability of forming pG4s in the transcriptome. It integrates three scoring systems: consecutive G over consecutive C score (cG/cC), G4Hunter (G4H) and G4 neuronal network (G4NN). For this study, sequences were considered positive if they obtained at least one score out of the three higher than the following thresholds: G4Hunter > 0.9, G4NN > 0.5, and cG/cC > 4.5. To specifically investigate APA events influencing mRNA length modulation, we excluded genes annotated with “Splicing” and focused our analysis on other events related to “shortening,” “lengthening,” and “both,” resulting in a total of 181 genes (Figure 2C). Under these stringent conditions, we identified 151 genes containing at least one pG4 near a PAC, thus confirming the prevalence of pG4 in 83% of PACs under RHPS4 treatment conditions. To streamline our focus on promising candidates, we refined our selection criteria by extracting APA events with a PDUI of more than 12%. Furthermore, we excluded cases involving introns and those with multiple exons containing PACs to enhance our research. As a result, we narrowed down our analysis to 15 genes, which encompassed a total of 27 pG4 structures (Supp Fig. 4).

**Figure 4:**
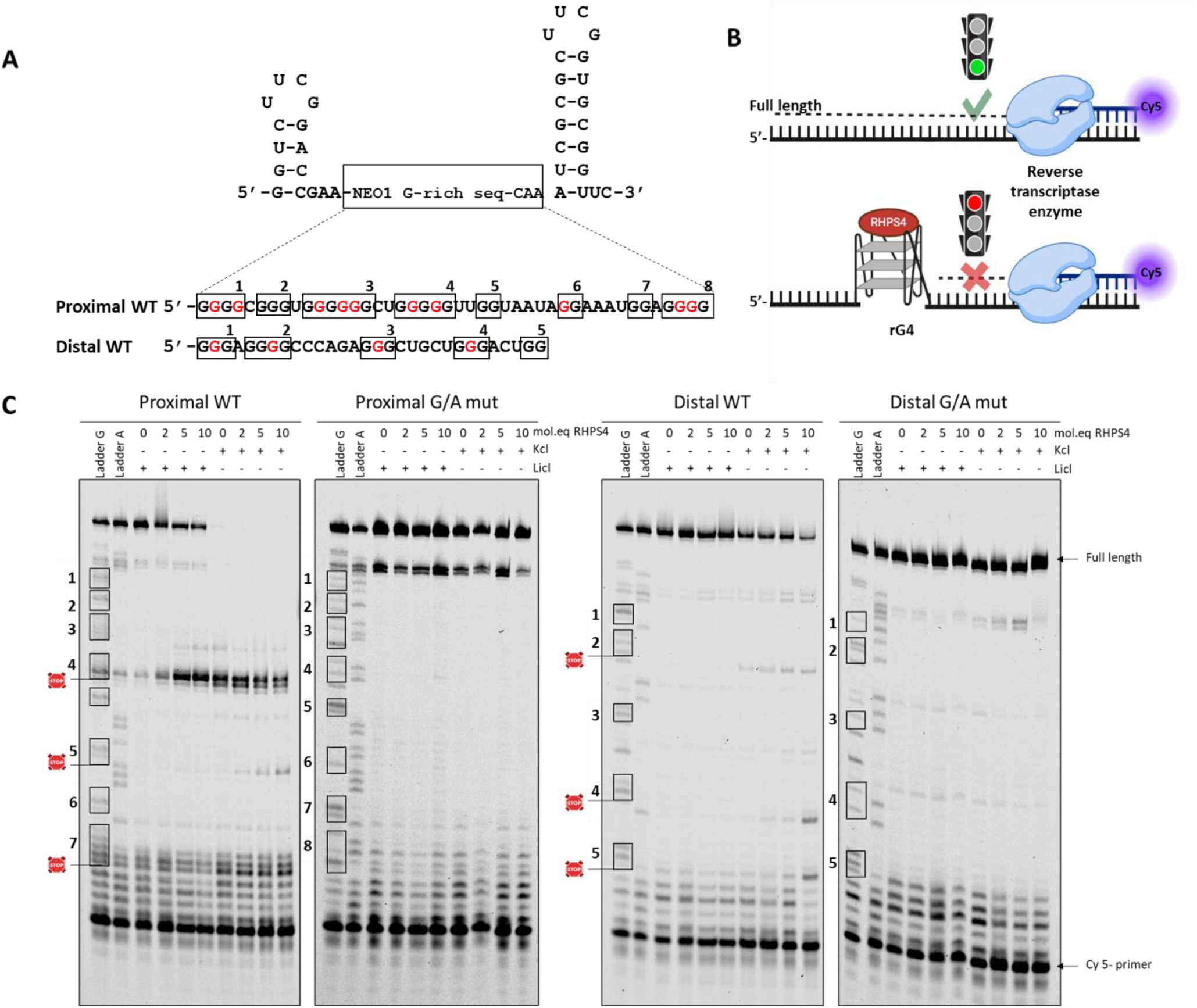
Assessing the stabilization and selectivity of RHPS4 on proximal and distal G4 of NEO1 3’UTR using reverse transcription stop assays (RTS) (A) Sequences of both proximal and distal G4 RNAs in their native (wild type) form are capable of folding the structure, while those with point mutations from G to A, highlighted in red, are unable to fold the G4 (G/Amut). These were inserted between the two hairpins. (B) Schematic diagram of the RTS reaction in the presence of RNA, with or without G4 stabilization by RHPS4 (C) RTS reactions were separated on 10% denaturing polyacrylamide gels. Each RNA was tested in the presence of increasing concentrations of the RHPS4 ligand (0, 2, 5, or 10 molar equivalents relative to the RNA concentration) under favorable conditions (K^+^) or unfavorable conditions (Li^+^) for G4 folding. Lanes labeled with “A” indicate the presence of ddTTP during the reaction, while those labeled with “G” were performed in the presence of ddCTP.

### pG4 near the proximal PAC of NEO1 3’UTR can form a G4

In the APA dataset, configurations with numerous PACs and identified pG4s can present challenges when selecting a candidate for the proof of concept, as mentioned earlier. Therefore, we endeavored to pinpoint an optimal scenario for the subsequent investigations. This process involved identifying a favorable spacing between our two PACs to facilitate the straightforward differentiation between the long and short mRNA forms, while ensuring the presence of no more than two pG4s. We chose to focus on the potential G4s (pG4) located near the PACs of NEO1. NEO1 protein is a single-pass transmembrane receptor belonging to the immunoglobulin (Ig) superfamily ^22^, playing essential roles in various cellular processes such as cell motility and adhesion (e.g., axon guidance and vascular development)^23,24^ as well as in survival and differentiation.^25^ In addition to being in a situation easy to study, membrane proteins (like NEO1) are highly cell-type specific, with differences in 3′-UTR length playing a significant role in regulation. They are typically regulated by interactions between their 3′-UTRs and RNA-binding proteins (RBPs), which serve to recruit effector proteins to the translation site and could potentially be disrupted by APA-shortened 3′UTRs.^26,27^ Visual representation of reads obtained for the 3’UTR of NEO1 and the nucleotide sequence revealed four PolyA cleavage sites through DPAC analysis: PAC1 and PAC2 near the stop codon (proximal region), and PAC3 and PAC4 in a distal area (highlighted in black in Fig. 3A) suggesting the production of two distinct mature RNAs with varying lengths of 3’UTR. Unexpectedly, treatment with RHPS4 prompted a notable preference for the distal polyA site selection, with a PDUI of 55% compared to untreated cells (PDUI = 37%), marking an 18% increase of long RNA form (Fig. 3A). Several *cis*-elements involved in the APA mechanism have also been identified: the consensus UGUA motif known to be recognized by the CFI complex (highlighted in blue in Fig. 3B),^28^ four specific six-nucleotide PAS sequences (highlighted in gray), typically located 10–40 nucleotides upstream of the polyA cleavage sites. Among the 18 PAS curated in the human genome, the distribution of each PAS identified in NEO1 has been determined through motif analysis conducted across four human databases.^29^ Additionally, a U/GU-rich downstream element (DSE) is located within approximately 40 nucleotides downstream from the cleavage sites (highlighted in orange in Fig. 3B). Furthermore, as previously mentioned, the G4RNA screener identified two potential rG4s (represented in red in Fig. 3B) near these PACs, solely based on the nucleotide sequence of the RNA. The first, named proximal G4, lies 32 nucleotides downstream of PAC2, while the second, termed distal G4, is positioned 59 nucleotides upstream of PAC3 (Fig. 3B). The proximal G4 shows a significantly elevated probability of forming a G-quadruplex, with scores of 2.2 for G4 Hunter, 0.99 for G4NN, and a cG/cC score of 77. Regarding the distal G4, it also demonstrates relatively high scores (G4 Hunter: 1.2, G4NN: 0.8, and cG/cC: 4.3), suggesting a propensity for G4 formation, albeit to a lesser extent (Fig. 3B).

To confirm their actual folding into G4 structures, fluorescence assays were conducted using the sequences of the two RNAs transcribed *in vitro*, corresponding to the proximal G4 and distal G4. We ensured to include nucleotides upstream and downstream (15-20 nt) to maintain the natural context of these structures, as neighboring sequences could impact the folding of a pG4.^30^ Additionally, we used the light-up probe, N-Methyl-Mesoporphyrin IX (NMM), which is known to emit fluorescence upon binding to a native rG4 structure.^31^ As anticipated, when binding to the proximal rG4, NMM exhibits fluorescence intensities 40 times greater than NMM alone (F/F0) which is comparable to characterized rG4.^14,32^ This finding remains consistent in the presence of both K^+^ (i.e. rG4 favorable folding conditions) and Li^+^ (rG4 unfavorable folding conditions), indicating the remarkably high prevalence of this rG4 structure. However, the distal G4 alone does not exhibit the ability to enhance fluorescence intensity, irrespective of the salt conditions in the solution (Fig. 3C). Not unexpected considering the low score achieved by the G4 RNA screener.

### RHPS4 stabilizes NEO1 3’UTR distal and proximal rG4-Structures

mRNA NEO1 harbors two G-rich sequences capable of forming natural G4 structures, as identified by G4RNA. The proximal G4 has been confirmed to form a G4 *in vitro* without requiring a ligand, while the distal rG4 structure confirmation may necessitate RHPS4. Considering the significant change in APA observed after RHPS4 treatment, both sequences may function as APA switches. Therefore, we aimed to determine if RHPS4 could enhance the formation of distal and proximal rG4 structures. To achieve this, the Reverse Transcriptase Stop Assays (RTS-assay) were conducted on both wild-type sequences (WT) and G/A-mutants designed to prevent rG4 formation by substituting certain Guanines (G) with Adenines (A). RTS-assay was employed to investigate the control of rG4 formation with a G4-ligand during reverse transcription.^13,14^ Essentially, the reverse transcriptase enzyme halts or slows down when encountering a stable structure such as a rG4. Consequently, it becomes feasible to detect this interruption using a Cy5-labeled antisense primer by separating the resulting single-stranded DNA on a denaturing gel (Fig. 4A). All RNAs (WT-proximal, WT-distal, GA-proximal and GA-distal) are embedded between a 5’ hairpin that serves as internal RTS control, and a 3’ hairpin for primer binding (Fig. 4B). RTS-assays were carried out in the presence of increasing concentrations of RHPS4 (0 to 10 mol.equiv/RNAs) under Li^+^ or K^+^ conditions (150 mM). For the WT-proximal G4, the band pattern immediately reveals stops from the first nucleotides corresponding to the last stretch of Gs (see box #8 in Fig. 4C) and a second stop at box #4 in all conditions (with or without RHPS4, in the presence of both Li+ and K+) validating the prevalence of the proximal G4 as demonstrated by the NMM assay. Conversely, compared to the GA-proximal, which shows no premature stops even in the presence of 10 eq RHPS4 in K^+^ conditions, confirming the total absence of G4-forming structures in the GA proximal mutant.

In the presence of Li+, there is a notable increase in premature stops following the addition of RHPS4 with the WT-proximal. However, the polymerase can still complete polymerization until the end, extending up to the 5’ hairpin, indicating incomplete formation of rG4 at this site (under unfavorable salt conditions). Nevertheless, in the K+ conditions, the stop profile remains similar, with only the addition of a third stop at box #6. The presence of RHPS4 does not seem to affect the intensity of the bands, and the reverse transcriptase fails to complete polymerization, highlighting the significant stability of the structure.

As mentioned, the distal G4 presents an interesting scenario. Despite the scores obtained from the *in silico* analysis of its sequence indicating the presence of a G4 structure, the results in solution with NMM suggest the opposite, even in the presence of K^+^. Therefore, the assays conducted on the WT sequence under Li^+^ conditions validate the fluorescence observations, indicating no formation of G4 structures. However, in the presence of K^+^, a faint band is observed at the level of Box #2, and the addition of RHPS4 intensifies this band while also causing premature stops to appear at the level of Box #4 and #5 with 10 eq RHPS4, indicating the formation of a weak distal rG4 under these specific conditions (K^+^ + RHPS4).

#### Mutations that impair distal G4 folding can alter APA of NEO1

Following confirmation of rG4s formation, we explored the individual roles of each rG4s in APA in cells. Briefly, the full-length 3′UTR, as well as the GA mutant versions, of NEO1 gene were cloned downstream of the open reading frame (ORF) of GFP in the pEGFP-C1 vector. This procedure included the removal of the SV40 polyA signal and the synthetic polyadenylation site already present in the vector. In total, four constructs were created through mutagenesis: one containing the WT NEO1 sequence with both proximal and distal G4 structures, another unable to fold the proximal G4 (GA proximal mutant), a third unable to fold the distal G4 (GA distal mutant), and a fourth with mutations preventing the formation of any G4 structure (double mutant; Fig. 5A). The transfection of plasmids and subsequent isolation of total RNA allowed to evaluate the abundance of long 3’UTR (GFP-NOE1 LU at 3.2 kb) and short isoforms (GFP-NEO1-SU at 1.5 kb) production 24 h post-transfection using Northern blot analysis, based on the mentioned constructs (Fig. 5B). Averaging data from three biological replicates, we calculated the PDUI for each construct using the formula (LU/(LU+SU): Fig. 5C and D). In our raw results, we found the PDUI for the wild-type (WT) 3’UTR sequence to be 0.22 of LU, which falls within the same range as the PDUI obtained through our DPAC analysis of the endogenous NEO1 gene (36% of LU in untreated cells). This suggests that the long isoform is unfavorable in both instances, indicating a consistent and reliable pattern of alternative polyadenylation for NEO1. Surprisingly, the mutation of the proximal G4 does not drastically change the PDUI (0.25), despite appearing to be the most formed in NMM and stabilized in RTS. Conversely, mutation of the distal G4 led to a significative reduction in the production of the long form (PDUI = 0.11). The double mutant exhibited an intermediate level at 0.15, similar to the distal mutant. While a significant direct effect on the APA has been observed, specifically on the PAS, the guanines implicated in the G4 folding found by RTS and NMM, both for the distal and the proximal, do not correspond to the conventional recognition sites (PAC, PAS, DSE). Overall, the distal G-quadruplex formation appears to have a direct impact on the polyA site that the RNA processing machinery selects.

**Figure 5.**
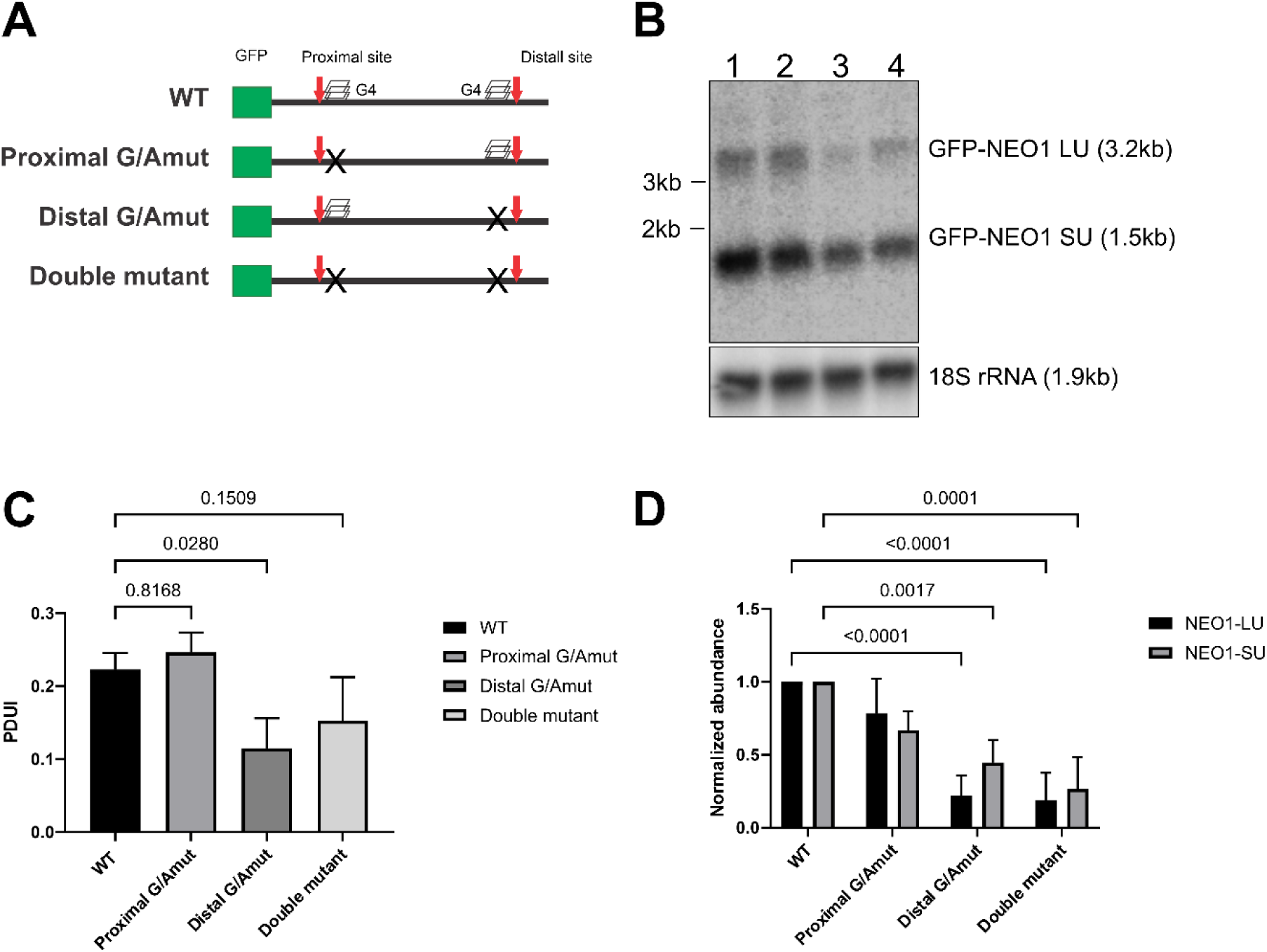
Mutations that prevent the formation of the distal G4 decrease the use of the distal polyadenylation site. (A) Graphical representation of RNAs transcribed after transfection of plasmids into HEK-293T cells. The 3’UTR of NEO1 is fused to the GFP sequence. In the diagram, the position of the proximal and distal poly(A) sites is indicated by red arrows while the G4 sequences are represented by stacked squares. When these are mutated, they are represented by Xs. (B) Typical Northern blot allowing the detection of both long (GFP-NEO1 LU) and short (GFP-NEO1 SU) isoforms using a GFP probe after a 24 h transfection. The loading control used here is the 18S RNA which can be seen under the membrane. Lane 1; WT, Lane 2; Proximal G/Amut, Lane 3; Distal G/Amut, Lane 4; Double mutant. (C) Intensity of GFP-NEO1 SU and LU from Northern blotting experiments was determined using Image J. Then PDUI (LU/(LU+SU)) was calculated. The experiments were repeated three times with each of the constructions being transfected once. The statistical analysis performed is a one-way ANOVA with Dunnett’s multiple comparison of each tested constructions performed vs WT. Adjusted p values are shown for each comparison. Error bars are standard deviation. (D) Intensity of GFP-NEO1 SU and LU from Northern blotting experiments was determines as in (C) except the intensities were normalized against 18S RNA (for loading control) and WT isoforms (for abundance). The experiments were repeated three times with each of the constructions being transfected once. The statistical analysis performed is a two-way ANOVA with Sidak’s multiple comparison of each tested constructions performed vs their respective WT. Adjusted p values are shown for each comparison. Error bars are standard deviation.

## Discussion

The expansive dataset described in this study demonstrates that rG4 is a prevalent feature of APA within the RNA landscape, as evidenced by its identification using a classic pharmacological probe targeted against the well-characterised rG4 motif (RHPS4). This G4-ligand has the potential to stabilize multiple types of G-quadruplexes and exert a multifaceted regulatory effect on various G4s, complicating the analysis of their regulatory role in G4-APA. However, through DPAC-seq analysis, we thoroughly demonstrate a correlation between the formation of rG4s around APA sites and significant changes observed when cells are exposed to RHPS4 treatment. Among the significant events, we identified the transcript of NEO-1 as an ideal case: the formation of 2 rG4s (so called proximal and distal) around 2 PACs (respectively from 32 and 59 nt) have been reported by an efficient bioinformatic pipeline. Upon stabilizing the G4 of NEO1 with RHPS4 in cells, we observe an increase in PDUI (from 0.36 to 0.55 and an elevation in the abundance of NEO1 transcript (refer to data analyzed with DPAC and DSeq2, respectively). Additionally, functional assays conducted with Northern blot fusion GFP-3’UTR NEO1 experiments highlights the significance of the distal rG4. Specifically, when the distal rG4 is mutated (rather than the proximal one), a decrease in PDUI (from 0.22 to 0.11) and a reduction in NEO1 expression abundance are observed. This decrease does not seem to be due to a loss of stability in the long isoform, as demonstrated by the mRNA stability assay conducted using transcription inhibition by Actinomycin D (see Supp Fig. 5). In all scenarios, the anticipated trend persists: the short isoform (SU) remains more stable compared to the long (LU). Nevertheless, in the context of the distal G/A-mut, this stability difference appears to mirror that observed in the wild-type. We also considered the potential impact of rG4 formation within the DNA on upstream transcription processes. Biophysical studies (including CD, NMM, and RTS-stop assays) generally indicate minimal to negligible formation of distal DNA G4 structures, as well as lower levels of proximal rG4 structuration compared to G4 RNA in NMM, see Supp Fig. 6 and 7. Our results indicate that the addition of RHPS4 has a minor effect on both proximal and distal DNA G4 structures, as demonstrated by CD spectra and RTS-stop assay Nevertheless, it’s worth mentioning that while the rG4 proximal mutation remains unchanged (likely unresponsive to RHPS4 addition), the mutation of the distal rG4 presents a noticeable effect. Although key elements of PolyA (PAC, PAS) are close to the mutated sequences, the rG4s are not part of the identified PAS or PACS. This clearly highlights the effect of the rG4 structure, especially on the distal rG4 on APA, which aligns with expectations. Recent studies have highlighted the significance of the distal site over the proximal one, indicating a sequential regulation of APA. This involves the initial cotranscriptional selection of the distal PAS followed by the post-transcriptional utilization of the proximal PAS.^33^ Various RNA-binding proteins, including splicing and polyadenylation modulators, such as the heterogeneous nuclear ribonucleoprotein (hnRNP) family, are likely to bind rG4s. The recognition of rG4 by these RBPs is critical, with the localization of the rG4 playing a significant role. Our data indicates that the G4 near a proximal PAC enhances the usage of the corresponding proximal polyadenylation sites (pPAS). In the cases of LRP5 and FXR1, among others, the rG4 is situated downstream of the pPAS and upstream of the distal PAS. Overall, this spatial arrangement appears to preclude the use of the distally located site causing a decrease of longer isoforms, as it is also observed for NEO1. This could enable a more precise control of the regulation of polyA sites by the formation of synthetic G-quadruplexes, and thereby allowing for transcripts with higher stability to be produced, while at the same time affecting preferential protein isoform production. Thus, this strategy may facilitate a shift in our view of secondary structures from adverse entities to powerful modulators of the proteome, impacting on all areas where these molecules have therapeutic and biotechnological potential.

## Author Contributions

F.B, P.L and J.P.P. conceptualized this study. F.B, P.L and M.A.T performed the experiments. FB, P.L. and M.A.T analyzed the data. P.L, F.B and J.P.P. wrote and revised the manuscript.

## Acknowledgments

We would like to thank Anais Vannutelli for her invaluable assistance and insightful discussions during the bioinformatic analysis of the PolyA-click seq and the G4 identification.

## Materials and methods

All Oligonucleotides used in this study were purchased from Integrated DNA technology (IDT) and were stored at −20 °C as 100 μM stock solutions in deionized water. The actual concentration of these stock solutions was determined by spectrophotometry at 260 nm (table of oligonucleotides in supp Table 1). RHPS4 was purchased from Bio-techne and dissolved in water at a concentration of 10 mM prior to storage. These stock solutions were then diluted with deionized water to the appropriate concentration for the experiment.

### Dose–response of RHPS4

Human embryonic kidney HEK-293T cells cultured in Dulbecco’s modified Eagle’s medium (Wisent) supplemented with 10% fetal bovine serum (Wisent) were seeded at 5000/well in a 96-well flat bottom plate. The day after, cells were treated with a series of RHPS4 (#5311, Bio-Techne) concentrations made from serial dilutions. Following an incubation time of 72 h, cell proliferation assays were performed using MTT assay kit (Abcam) according to manufacturer’s recommendation. Three biological replicates were carried out to generate the dose-response curve from which the inhibitory concentrations (IC) were calculated (Supp. Fig. 1).

### Poly(A) click seq

HEK-293T cells were seeded at 5×10^5^ in a 10 cm petri dish. The next morning, RHPS4 was added to a final concentration of 1.5 µM and incubated for 72 h at 37 °C in a humidified, 5% CO_2_ atmosphere-controlled incubator. In parallel, a non treated petri dish was also incubated for the same time period. Total RNA was then isolated using Qiazol, according to the manufacturer. Following suspension in high pure water, the total RNA was further purified using RNAspin Mini isolation kit (Cytiva) that includes on-column DNAse I digestion according to manufacturer’s protocol. Total RNA was eluted, concentration determined by spectrometry at 260 nm using a NanoDrop spectrophotometer (Thermo Fisher Scientific, Mississauga, ON) and quality assessed following separation on agarose gel. Three biological replicates were carried out before sending these samples to the company Clickseq Technologies LLC (Davis, CA, USA) in order to analyze the changes in polyadenylation by Poly(A) click seq. Library preparation and sequencing were performed as described previously. ^16^ Data analysis pipeline was performed using DPAC software for poly(A) changes, and DESeq2 for gene expression variations as described previously. ^34^ Single-end run (150 bases) was performed on an Illumina NextSeq 550. Reads were mapped on Homo Sapiens genome assembly (hg19). The results were returned to us in an excel file. Using the program DPAC, poly(A)-Sites (PASs) are defined as the exact nucleotide of 3’UTR to poly(A)-tail junction identified with at least 5 mapped reads and with at least 25 A in the read. All provided samples are used to generate a project specific database of Poly(A)-Clusters (PACs), which is subsequently used for differential gene and alternative polyadenylation analysis. PACs are defined as groupings of poly(A)-sites that are found within 10 nucleotides of one another. Gene counts for each sample are generated using DPAC. Using the newly generated annotated PAC database, the 3’ end of all mapped reads overlapping PACs within a gene contribute to that gene count. The count table is input into DESeq2 for differential gene expression analysis. Differential gene expression is defined as >1.5-fold change with p-adjust <0.1. If there are more than one Poly(A)-Clusters in a 3’UTR, and one or more of them are ‘significantly’ regulated following the addition of RHPS4, then we can say that there is lengthening or shortening APA (according to DESeq2 and DPAC), if there is a change in any 3’ end alternative isoform by greater than 10% (p-adjust <0.1). Moreover, for each significantly regulated event, a differential percentage of distal poly(A) site usage (ΔPDU) il also generated. The latter considers the length of the 3’UTRs in each isoform and weights the contribution of a PAC based upon its genetic distance from the most proximal PAC (no statistical test involved).

### Bioinformatics analyses for the presence of potential G-quadruplexes near PACs

For each significant event detected, depending on the location of the PACs on the mRNA, there are different types of poly(A) events (see Fig. 1). To simplify the analysis, we restricted the search for events that could involve G-quadruplexes to that called APA-exon where the PACs involved are located strictly in the last exon, i.e., the 3’UTR. Thus, for each PAC involved in a significant poly(A) change following RHPS4 treatment (called by DPAC “lengthening”, “shortening” or “both”), we used their chromosomal location to search for the presence of G-quadruplexes up to 100 nt upstream and downstream. In order to do that, we used G4 RNA screener that contains three scoring systems to describe the likelihood of G4 observation: cG/cC score, G4Hunter (G4H) score and G4NN score. For this study, sequences considered positive imply obtaining at least one score out of three higher than the following limits: G4H > 0.9, G4NN > 0.5, cG/cC > 4.5. Subsequently, for each PAC positive for the presence of a potential G4 sequence, we added another level of control by removing all situations for which the ΔPDU is less than 0.12. Finally, some hits were manually cleaned by removing false positive sequences generated by the cG/cC score alone.

### Gene ontology

The Gene Ontology analysis was performed on the list of significantly up-regulated and down-regulated genes regulated by RHPS4 addition. The GO term analysis was conducted using the website https://geneontology.org/, in accordance with the GO Ontology database released 2023-05-10, DOI: 10.5281/zenodo.7942786

### Fluorescence assays

RNA molecules were synthesized by *in vitro* transcription using purified T7 RNA polymerase. Briefly, two oligodeoxynucleotides (2 μM each, Life Technologies) were annealed together through complementary regions, and then purified Pfu DNA polymerase was used in PCR reactions in the presence of 5% DMSO to fill in the gaps. The resulting duplex DNA products were then ethanol-precipitated, washed with 70% ethanol, and dissolved in ultrapure water. Run-off transcriptions were then performed in a final volume of 100 µL using purified T7 RNA polymerase in the presence pyrophosphatase (0.01U, Roche Diagnostics) and 5 mM concentrations of each nucleotide triphosphate in a buffer containing 80 mM HEPES-KOH (pH 7.5), 24 mM MgCl_2_, 40 mM dithiothreitol (DTT), and 2 mM spermidine. The reactions were incubated for 2 h at 37 °C followed by DNase RQ1 (Promega, Madison, WI) treatment (2 Units) for 30 min. at 37 °C. The RNA was then purified by phenol:chloroform extraction and recovered by ethanol precipitation. The RNA products were then fractionated by denaturing (8 M urea) 8% polyacrylamide gel electrophoresis. The RNAs in the gels were detected by ultraviolet shadowing, and the bands corresponding to the appropriate sizes were excised from the gels. The transcripts were then eluted overnight at room temperature in a buffer containing 1 mM EDTA, 0.1% sodium dodecyl sulfate and 0.5 M lithium acetate, ethanol-precipitated, dried, and dissolved in water, and their concentrations were then determined by spectrometry at 260 nm using a NanoDrop spectrophotometer (Thermo Fisher Scientific, Mississauga, ON). In order to evaluate G4 formation, *in vitro* synthesized RNAs (200pmol) were added to a folding buffer containing (20 mM Li-cacodylate pH 7.5, 20 mM MgCl_2_ and 100 mM of either LiCl or KCl) and were then heated at 70°C for 5 min before being slowly cooled to room temperature over a period of 1 h. The reactions were then completed to 100 µL with a buffer containing 20mM Li-cacodylate pH 7.5, 20 mM MgCl_2_ and 100 mM of either LiCl or KCl). Next, 2.5 eq/RNA of N-Methyl-Mesoporphyrin IX (NMM) (Frontier Scientific Inc., Logan, Utah) was added, and the reactions incubated in the dark for 5 min at room temperature in a 10 mm quartz cuvette. The fluorescence intensity was then monitored using a Hitachi F-2500 fluorescence spectrophotometer with an excitation wavelength of 399 nm and the emission spectra were recorded between 500 and 650 nm. The fluorescence at 605 nm was used for quantification. All experiments were performed in duplicate.

### Reverse transcriptase stops assays (RTS)

RNAs to be tested in RTS are embedded between a 5’ hairpin that serves as internal RTS control, and a 3’ hairpin for primer binding (Fig. 5B)^35^. All reactions were performed in favorable conditions for G4 folding (KCl) or unfavourable conditions (LiCl). *In vitro* synthesized RNAs were generated as described earlier (see Florescence assays section), through ODNs annealed and gap filling using a T7 promoter ODN in a PCR reaction followed by *in vitro* transcription of the double-stranded DNA templates using T7 RNAP. The resulting RNAs were quantified and 1.5μL (3 μM) was added to 1 μL of 5 μM Cy5 fluorescently labelled DNA primer complementary to the 3’ hairpin. The reaction was completed by the addition of 5 μL of binding buffer (100 mM Tris-HCl pH 8.3, KCl 150 mM or LiCl 150 mM) and 1 μL of 0, 2, 5 or 10 molar-equivalents of RHPS4. The temperature of the mixture was then increased to 75°C for 3 min and kept at 37°C for 5 min in a thermocycler. At the beginning of the 37°C step, 0.5 μl of 10 mM dNTPs, 1 μl of 100mM DTT, 1 μl of 25mM MgCl_2_ and 1 μl of water, ddCTP or ddTTP was added. For each RNAs to be tested, ddCTP and ddTTP were added in reactions containing LiCl and not KCl to generate ladders. The 12 μL mixture was heated up to 45°C, and 0.5 μL (100 U) of purified MMuLV-reverse transcriptase was added to make up the 12.5 μl reactions. The reverse transcription was maintained at 45°C for 15 min, and then 0.5 μl of 2M NaOH was added at the end of the step. The temperature was immediately ramped up to 90°C for 10 min to inactivate the enzyme and degrade the RNA template. Finally, 15 μL of loading dye (98% formamide, 10 mM EDTA) was added and 6 μL was separated on a 10% polyacrylamide gel containing 8 M urea. The final images were generated using a GE Typhoon FLA 9000 fluorescence scanner (λex = 635 nm).

### Plasmids and cloning

Before cloning of the NEO1 3’UTR into pEGFP-C1 plasmid, we had to remove the synthetic SV40 polyadenylation signal sequence from position 1519-1640 in order to prevent any bias that could result from the unwanted competition between poly(A) sites from NEO1 3’UTR and SV40 poly(A) sequence. Plasmid pEGFP-C1 was used as a template in a PCR reaction using Q5 DNA polymerase (New England Biolabs) and oligonucleotides (5’-ACGCGTAAATTGTAAGCG-3’ and 5’-AACAACAACAATTGCATTCATTTTATG-3’) resulting in PCR fragment without region 1519-1640 of the plasmid. The PCR reaction was then purified using PCR purification kit (Bio Basic) according to manufacturer’s recommendation. After a KLD reaction, the resulting circularized plasmid (called pEGFP-C1 ΔSV40) was transformed into *E. coli* STBL3. Deletion of the synthetic SV40 poly(A) signal sequence was further confirmed by sequencing. A STOP codon was also introduced at the end of the GFP sequence. In this case, plasmid pEGFP-C1 ΔSV40 was used in a Quick-change reaction using a pair of complementary oligonucleotides. The resulting nicked-plasmid (called pEGFP-C1 ΔSV40-Stop) was transformed into *E. coli* STBL3. Introduction of the STOP codon (TAA) downstream of the GFP open reading frame was confirmed by sequencing. NEO1 3’UTR was isolated from genomic DNA isolated from cultured HEK-293T using DNeasy Blood and Tissue kit (Qiagen) according to manufacturer’s protocol. Purified genomic DNA (135 ng) was used as a template in a PCR reaction. And the 3’UTR of NEO1 (2 654 bp) was introduced into linearized pEGFP-C1 ΔSV40-Stop with *Xho* I and *BamH* I use the HiFi DNA Assembly Cloning Kit (New England Biolabs) according to manufacturer’s instructions. All of the mutants were created using the same protocol. Following the transformation of the ligation products into bacteria and the sequencing of the resulting plasmids confirmed the presence of the mutations.

### Northern blots

To generate GFP RNA probe (320 nt), DNA template for *in vitro* transcription was amplified from pEGFP-C1 with primers 5’-taatacgactcactataGGGTTACTTGTACAGCTCGTCCATG-3’ and 5’-CACAAGCTGGAGTACAACTAC-3’, where smallcap letters correspond to the T7 promoter sequence using purified Taq DNA polymerase in 100µL reaction. DpnI (1 μL) was added to the reaction to degrade original pEGFP-C1 template and incubated at 37°C for 1 h. Phenol-chloroform extraction followed by ethanol precipitation was performed. Pellet was dissolved in water and concentration measured with a Nanodrop spectrophotometer. 18S probe (80 nt) was generated using the antisense oligonucleotide 5’-CCTGGCGGAGCGCTGAGAAGACGGTCGAACTTGACTATCTAGAGGAAGTAAAAGTCGTAACAAGGTTTCCGTAGGTGAACCCtatagtgagtcgtatta-3’ and the sense T7 promoter as described previously for the GFP probe except, the PCR product was purified using PCR purification kit (Bio Basic). DNA eluted in pure water and concentration measured with a Nanodrop spectrophotometer. RNAs were synthesized using MaxiScript T7 kit (Invitrogen). Following transcription, probes were purified using MicroSpin G50 columns (Cytiva) and added to pre-hybridization buffer.

HEK-293T cells were seeded at 5×10^5^/well in a 6-wells plate. The day after, DNA plasmids (1 µg) containing NEO1 3’UTR-WT, NEO1 3’UTR-G/Amut proximal, NEO1 3’UTR-G/Amut distal and the double mutant fused to GFP were transfected with 3 µL of 1 mg/mL of polyethylenimine (PEI). Following an incubation time of 24 h, total RNA was prepared using 1mL of Qiazol (Qiagen) per well, essentially described in the previous section “Poly(a) click seq”. Again, concentration was determined by spectrometry at 260 nm using a NanoDrop spectrophotometer (Thermo Fisher Scientific, Mississauga, ON) and quality assessed following separation on agarose gel. To perform Northern blot analysis, 7.5 µg of total RNA was dissolved in 30 mM Tricine, 30 mM triethanolamine, 50% formamide, 0.4 M formaldehyde, 0.5 mM EDTA pH 8.0 and trace amount of bromophenol blue before being denatured at 70°C for 5 min. RNAs were separated on denaturing 1% agarose gel containing 30 mM Tricine, 30 mM triethanolamine and 0.4M formaldehyde at 6V/cm in migration buffer (30 mM Tricine, 30 mM triethanolamine). RNA was then transferred on Hybond-N+ membrane (Cytiva) 0/N by capillarity in SSC 20X (3 M NaCl and 0.3 M sodium citrate). After UV-crosslinking, the membrane was pre-hybridized in a buffer containing 50% formamide, SSC 5X, 1% SDS, Denhardt 5X and 100 μg/mL salmon sperm DNA for 4 h at 65°C. Internally ^32^P-radiolabelled RNA probe complementary to GFP sequence was then added and incubated O/N at 65°C. Three washes of 15 min each using 1X SSC, 0.1% SDS and one wash of 0.1X SSC, 0.1% SDS were performed at 65°C before exposition on a phosphoscreen and detection on a GE Typhoon FLA 9000 scanner. The membrane is used again for probing of the 18S RNA (for loading control) using the same steps as with the GFP probe. Image J software was used to measure band intensities and GraphPad Prism to generate histograms.

### Actinomycin assays

Stability tests in the presence of actinomycin D were carried out as described above under “Northern blots”. The difference is that 24h after transfection of the plasmids, actinomycin D dissolved in DMSO (final concentration of 6ug/mL) is added to each well, and incubated for times of 0-8h. At each time point, the RNA is isolated by Qiazol and analyzed by Northern blot as described above.

### Primer extension

DNA with a 3’ hairpin (750 nM) was incubated with a ^32^P radiolabelled reverse primer (750 nM) in a solution containing 20 mM Li-Cacodylate (pH 7.5) at 95 °C for 5 min, and the reactions were then cooled down at a rate of 1 °C/min to room temperature. The samples were then diluted to obtain final concentrations of 50 mM Tris (pH 7.6), 1 mM dNTPs, 1 mM MgCl2, 5 mM DTT, 100 mM KCl/LiCl and 675 nM ligand (diluting the RNA to 75 nM). The samples were then incubated at 37°C for 5 min. An in-house prepared Taq polymerase (2.5 U) was then added, and the reaction was further incubated at 37 °C for 30 min. The ladders were prepared in a PCR machine. Briefly, a solution containing 55 μM DNA 3’ hairpin, 55 μM of the radiolabeled primers, Thermopol buffer (NEB), 0.15 % Triton, 2U of Vent® (exo) DNA Polymerase (NEB), 54 μM ddGTP, 5.4 μM dGTP, 4.5 μM dATP and 15 μM of both dTTP and dCTP was incubated for 25 cycles of 1 min at 95 °C, 1 min at 52 °C and 1 min at 72 °C. The reactions were stopped by the addition of alkaline formamide loading dye (two volumes of 90 % formamide, 10 mM NaOH). The reactions were fractionated by electrophoresis on denaturing (8M urea) 10% polyacrylamide gels. The gels were dried, exposed on a phosphor screen, and visualized on a Typhoon FLA 9000 fluorescence scanner. All assays were performed in duplicate.

### CD analysis

The DNA samples were dissolved at a concentration of 4 μM in a solution containing 100 mM of either KCl or LiCl and 20 mM of Li-cacodylate (pH 7.5), with or without the presence of RHPS4 (10 eq). The samples were then heated at 90 °C for 5 minutes and slowly cooled down to room temperature. The experiments were performed in a quartz cell with a path length of 1 mm on a Jasco J-810 spectropolarimeter. Data were recorded from 350 to 210 nm at 20 °C. All CD spectra were smoothed (4 spectrum accumulations) and analyzed using Prism.

## SUPPLEMENTARY FIGURES

**Supp. Table 1:**
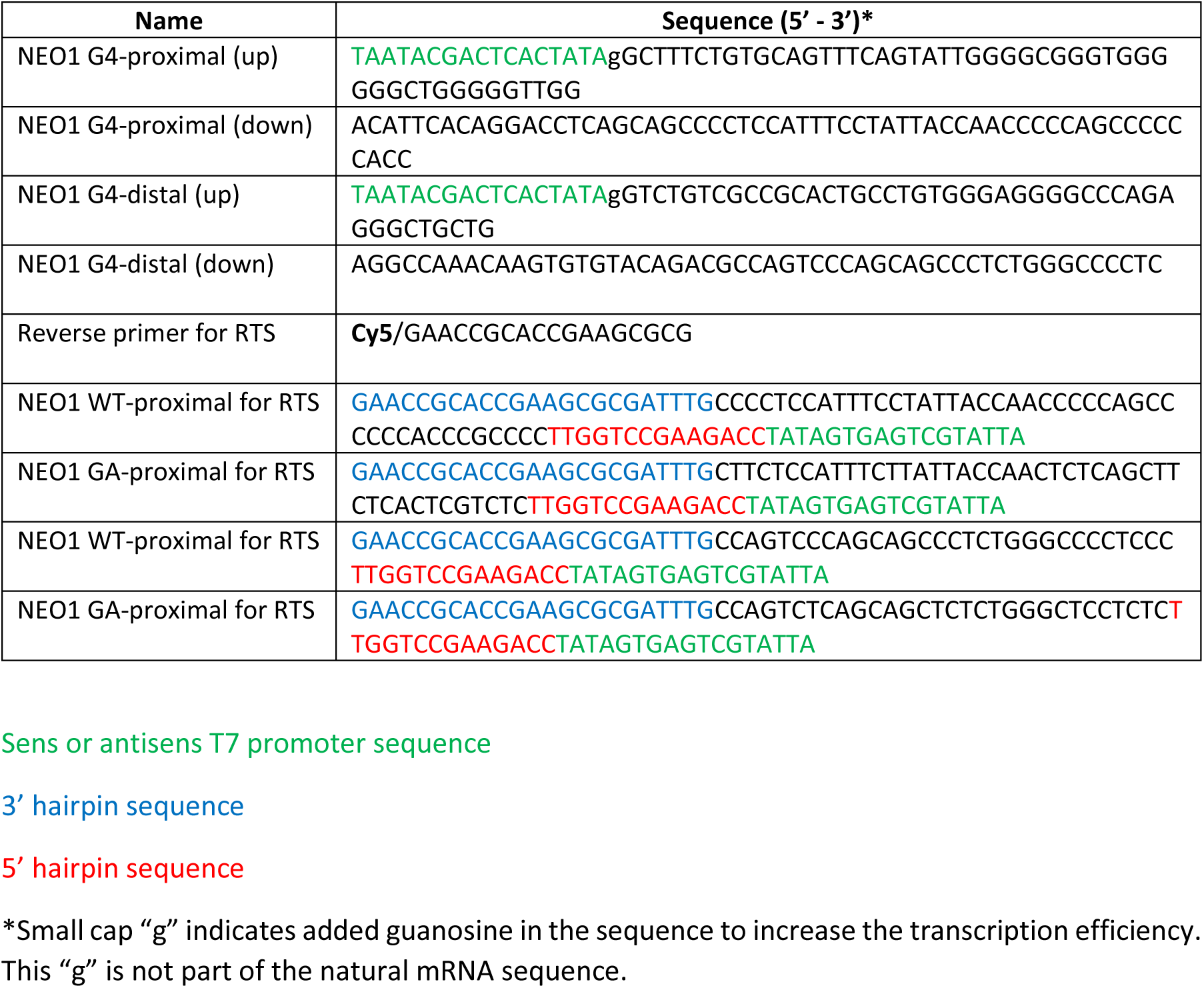
the oligonucleotide sequences.

**Supp. Fig 1:**
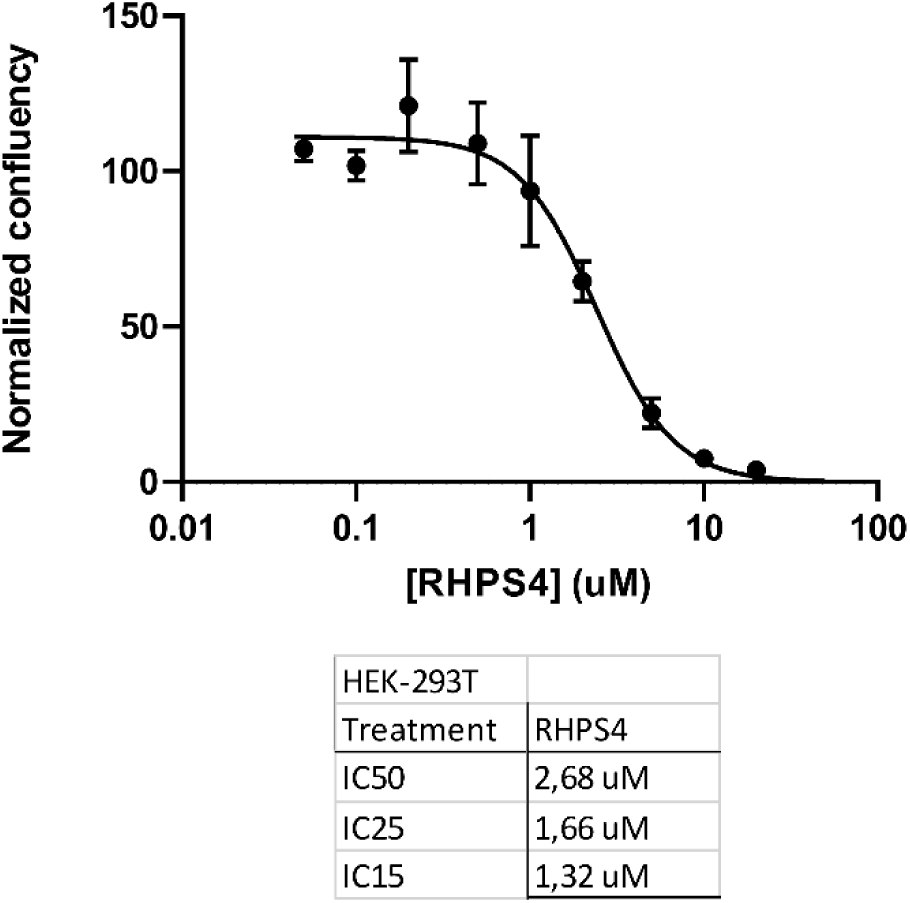
Dose–response of RHPS4 in HEK-293T. Dose response profile for RHPS4 in HEK-293T cells generated from MTT assays. Experiments were conducted with three biological replicates. The IC50, IC25 and IC15 are listed in the table below. Error bars represent standard error of the mean.

**Supp. Fig 2:**
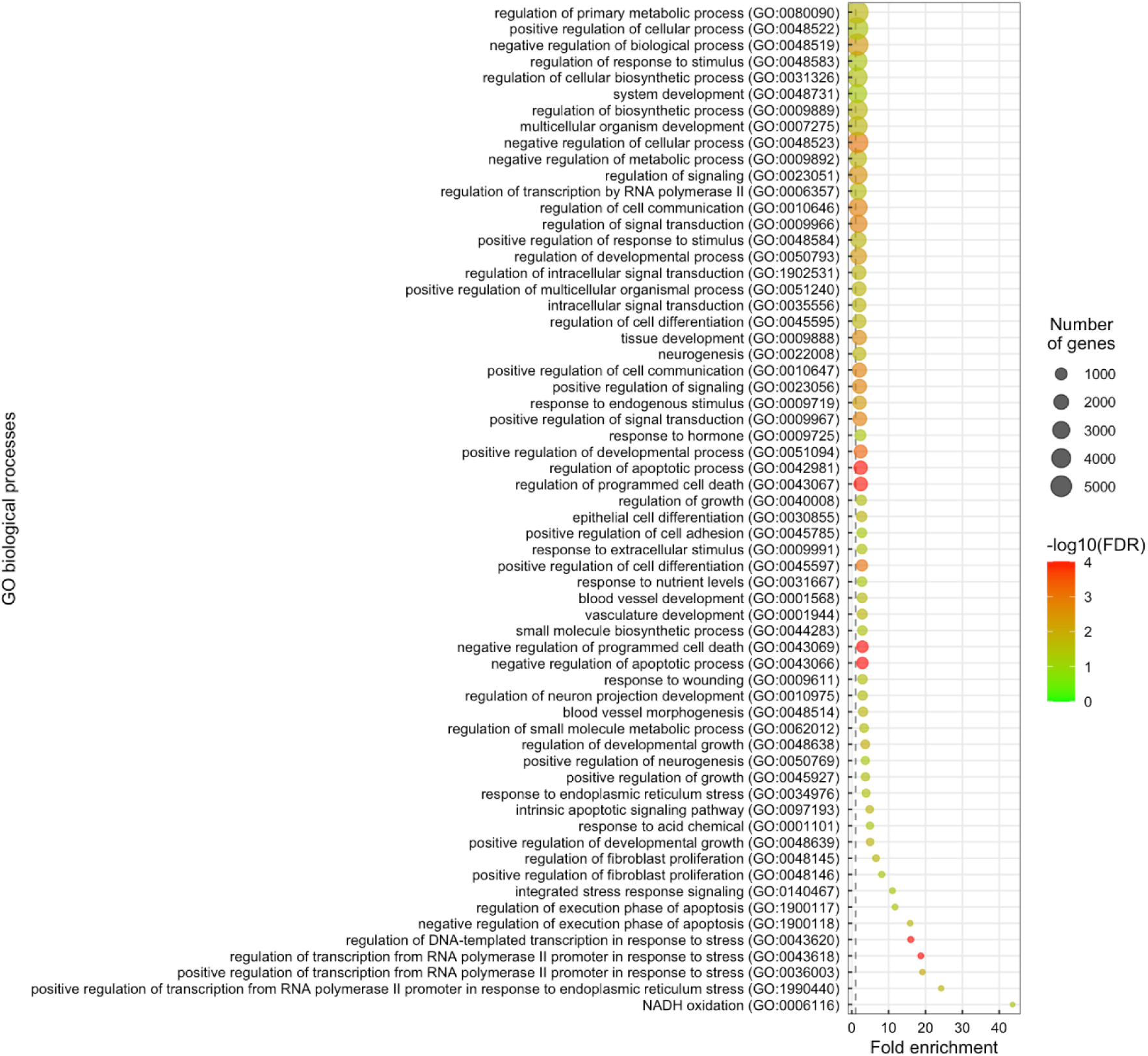
Gene ontology for genes that are up-regulated when cells are treated for 72 h with RHPS2.

**Supp. Fig 3:**
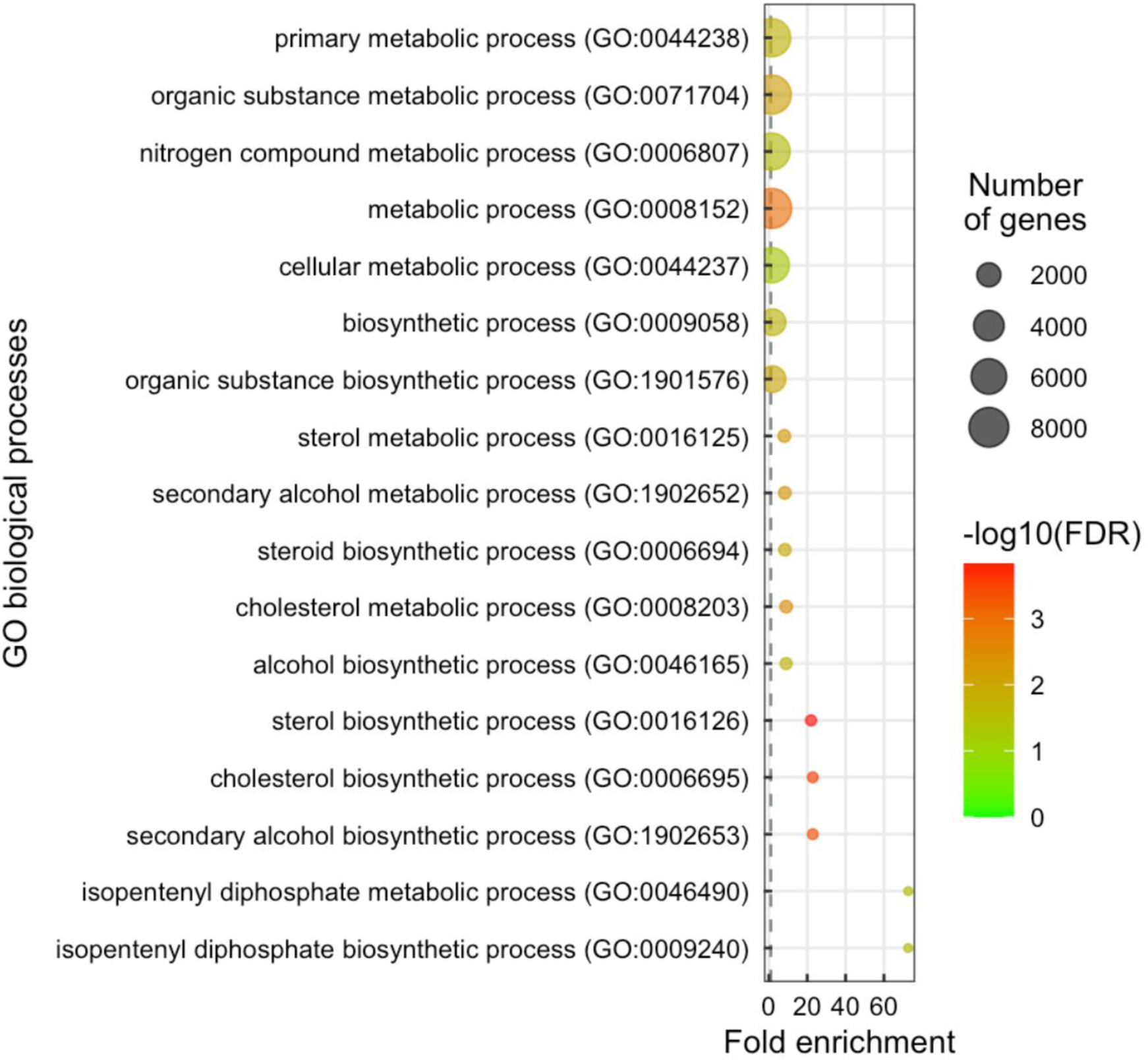
Gene ontology for genes that are downregulated when cells are treated for 72 h with RHPS4.

**Supp. Fig 4:**
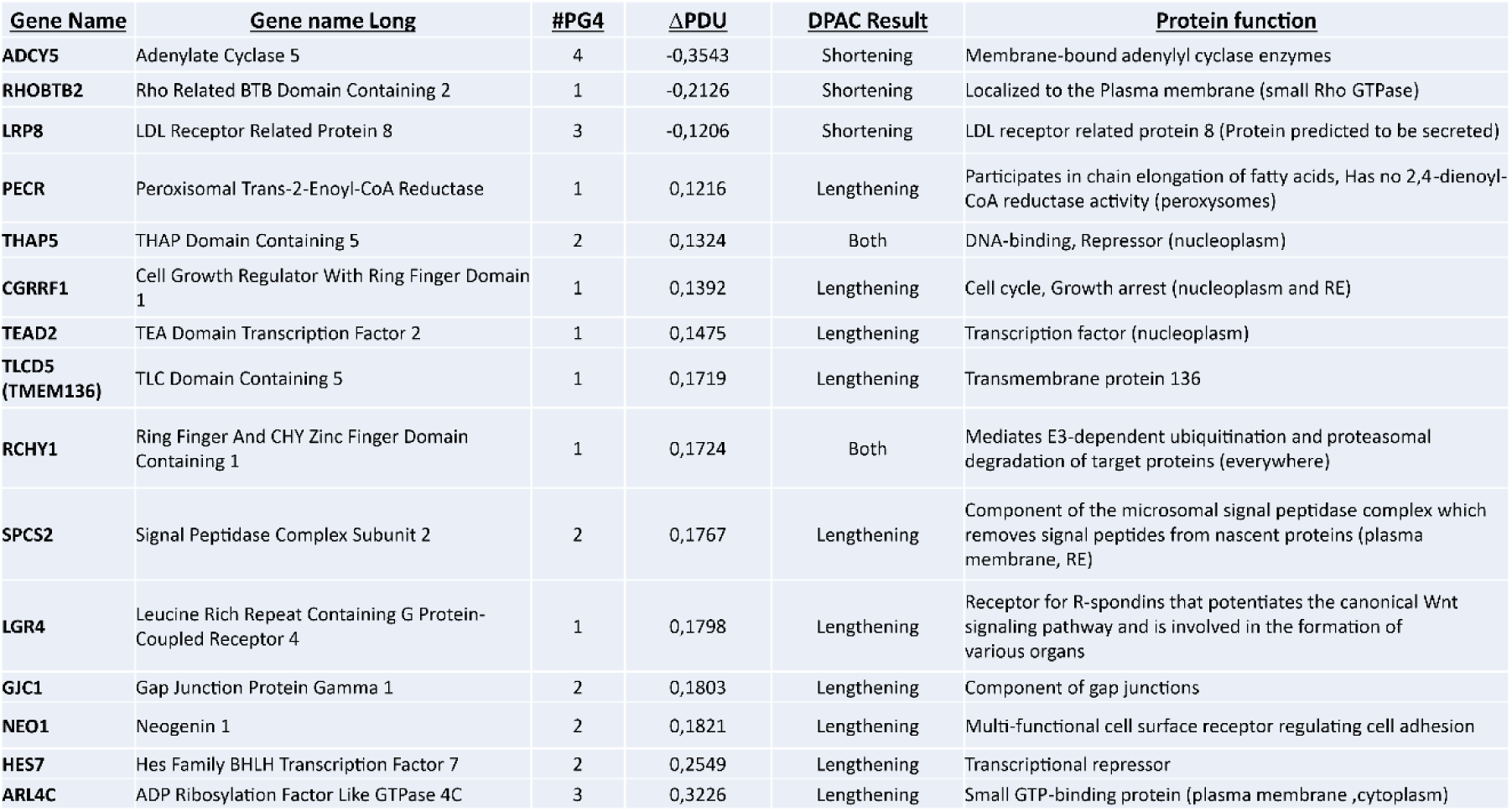
Candidate genes having at least one PG4 near a PAC. List of genes for which a significant variation in APA was recorded and containing at least one PG4 in the environment of PACs detected by DPAC. The table compiles for each gene, the number of PG4s involved nearby that can influence the APA, the ΔPDUI (long 3′UTR expression divided by long 3 ′UTR expression + short 3′UTR expression) as well as the result of the DPAC software and the function of the associated protein.

**Supp. Fig. 5:**
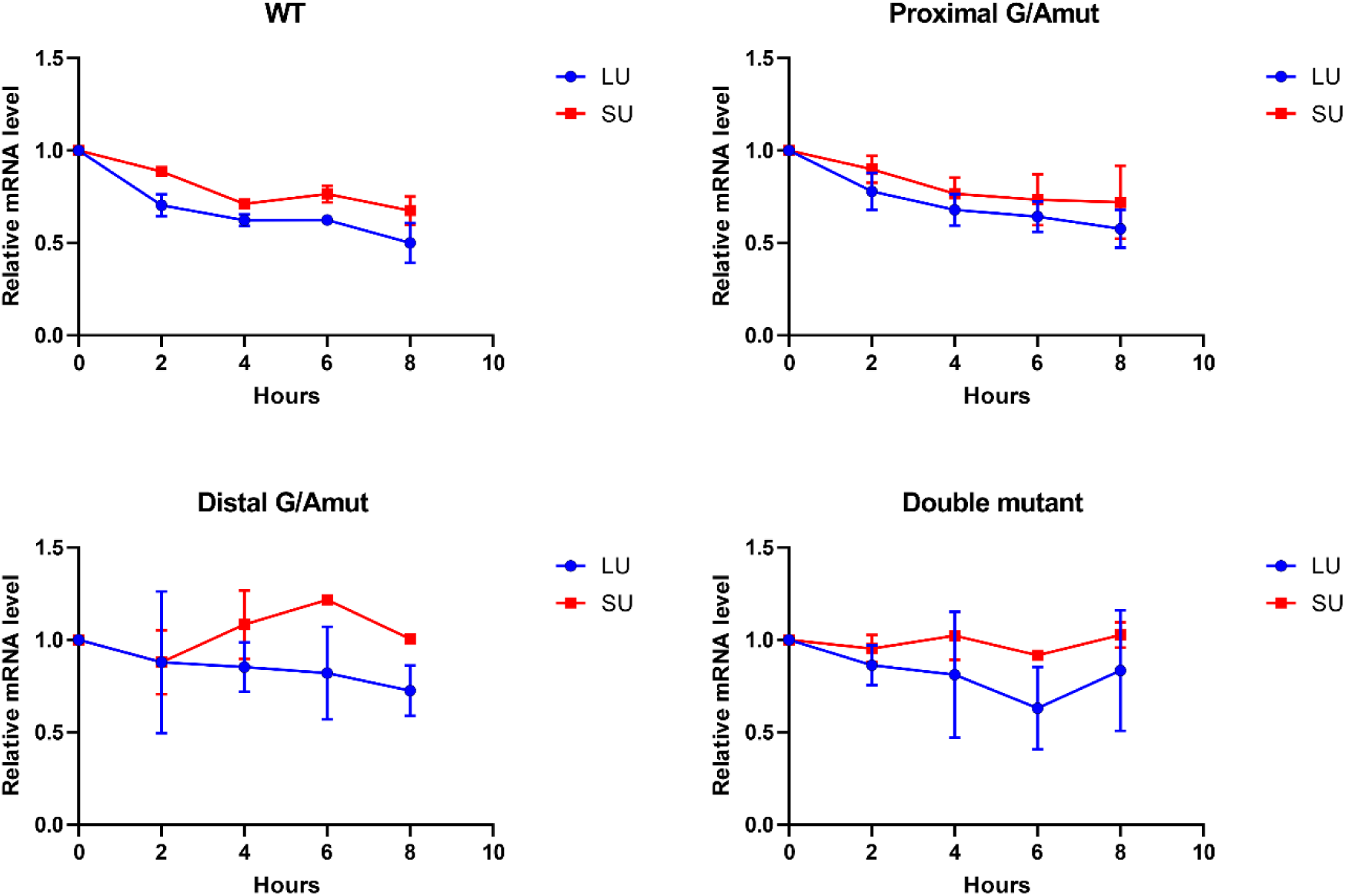
Stability assays with actinomycin D. Each plasmid construct containing the GFP-NEO1 3’UTR fusions and the various mutants (WT, proximal, distal and double mutant) was transfected and 24 h post-transfection, actinomycin D was added. After 0, 2, 4, 6 and 8 h of incubation, total RNA was isolated. The latter were separated on a denaturing agarose gel in order to carry out Northern blots with a GFP probe. The intensity of the bands (long (LU) and short 3’UTR (SU)) was determined using Image J software using 18S RNA as loading control. These graphs are the results of two biological replicates.

**Supp. Fig. 6:**
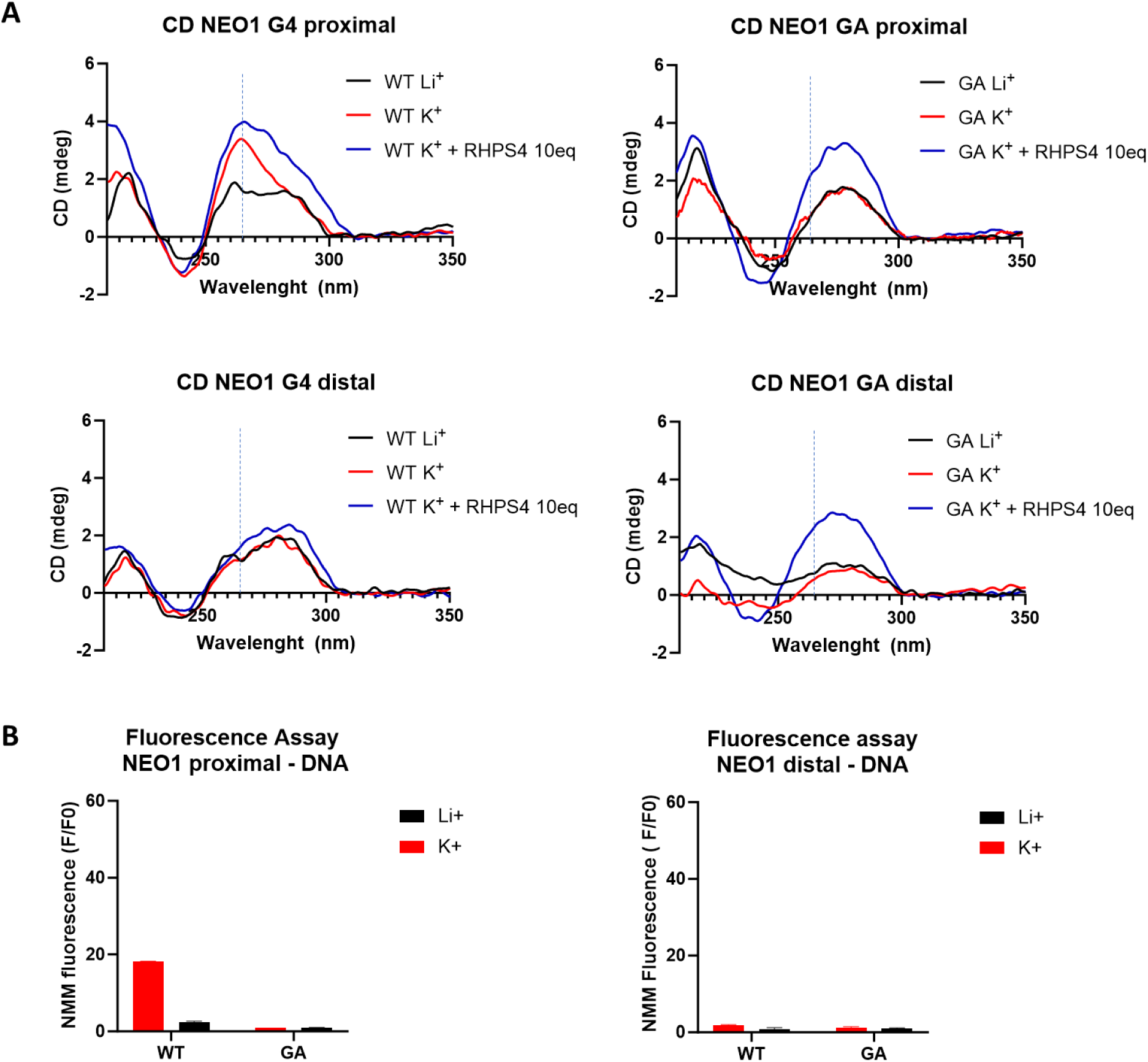
*In vitro* characterization of the DNA G4s corresponding to the proximal and distal G4 identified in NEO1 mRNA using fluorescent assay and circular dichroism. A) The circular dichroism spectrums were obtained in the presence of 4 μM of DNA and 100 mM of potassium with or without RHP4 (10 molecular equivalent). B) NMM fluorescence assays were performed in the presence of 2 μM of DNA, 4 μM of NMM and 100 mM of either KCl or LiCl at room temperature. All experiments were performed in duplicate and analysed by prism.

**Supp. Fig. 7:**
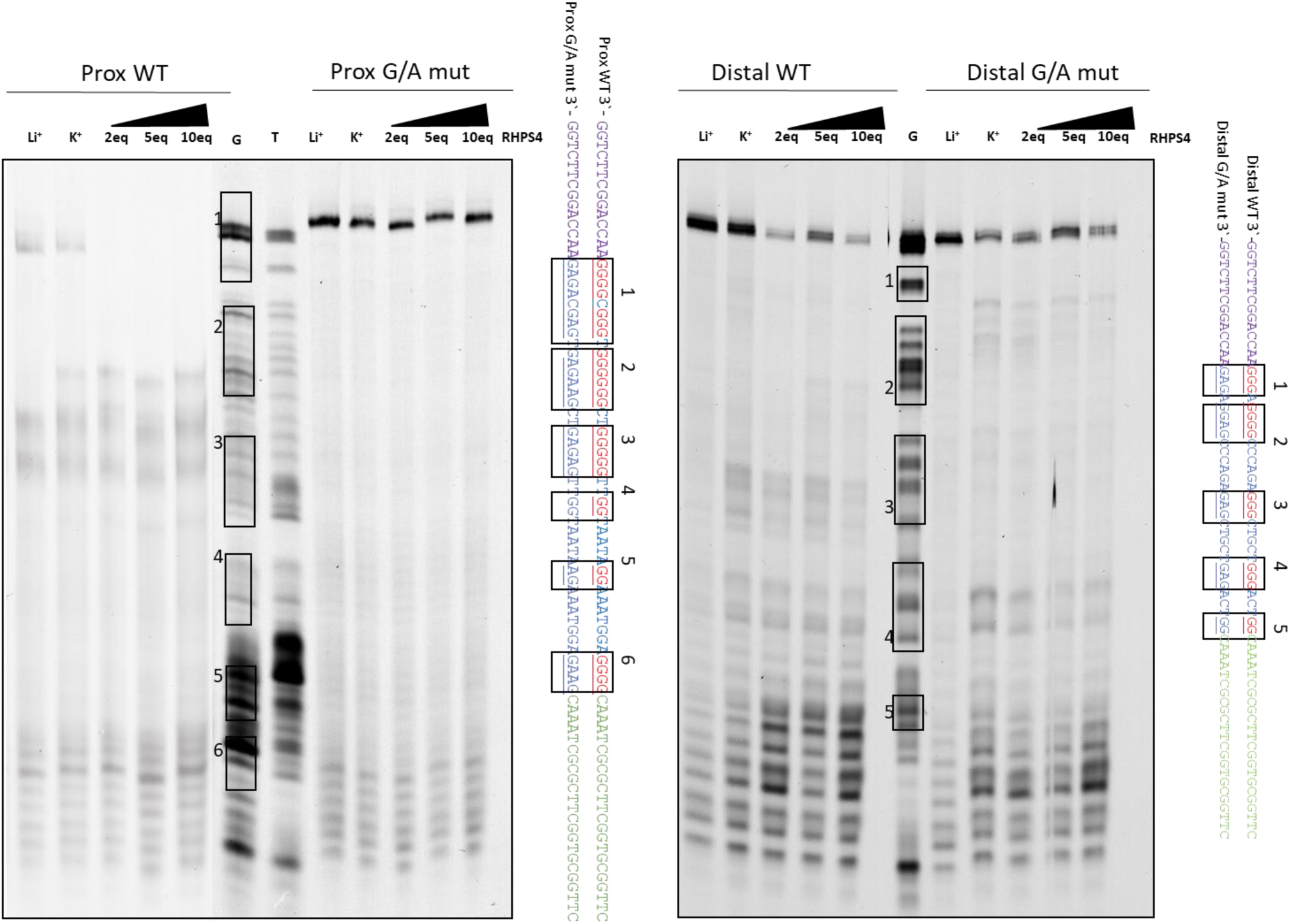
Evaluation of RHPS4-induced stabilization effects on DNA proximal and distal G4 structures using primer extension assays (PE). The DNA with a 3’ hairpin samples (750nM) were submitted to PE assays with an in-house prepared Taq polymerase (2.5 U) for 30 min at 37 °C. The reactions were performed in the presence of increased amount of RHPS4 (2 to 10 mol.eq), and a ^32^P primer in KCl buffer. The reactions were separated on denaturing 10% polyacrylamide gels. The red nucleotides represent those mutated in the G/A mutant.

